# Sequestrase chaperones protect against oxidative stress-induced protein aggregation and [*PSI^+^*] prion formation

**DOI:** 10.1101/2023.10.18.562867

**Authors:** Zorana Carter, Declan Creamer, Katerina Kouvidi, Chris M. Grant

**Affiliations:** Faculty of Biology, Medicine and Health, School of Biological Sciences, The University of Manchester, Michael Smith Building, Oxford Road, Manchester, M13 9PT, United Kingdom

## Abstract

Misfolded proteins are usually refolded to their functional conformations or degraded by quality control mechanisms. When misfolded proteins evade quality control, they can be sequestered to specific sites within cells to prevent the potential dysfunction and toxicity that arises from protein aggregation. Btn2 and Hsp42 are compartment-specific sequestrases that play key roles in the assembly of these deposition sites. Their exact intracellular functions and substrates are not well defined, particularly since no stress sensitivity has been reported in deletion mutants. We show here that Btn2 and Hsp42 are required for oxidant tolerance and act to sequestering misfolded proteins into defined PQC sites following ROS exposure. We have used the Sup35 translation termination factor as a model oxidized protein to show that protein aggregation is elevated and widespread in mutants lacking Btn2 and Hsp42. Oxidant-induced prion formation is also elevated in sequestrase mutants consistent with the idea that Btn2 and Hsp42 function to sequester oxidatively damaged Sup35, thus preventing templating to form its heritable prion form. Taken together, our data identify protein sequestration as key antioxidant defence mechanism that functions to mitigate the damaging consequences of protein oxidation-induced aggregation.

## Introduction

Proteostasis is maintained by an arsenal of molecular chaperones that are able to detect non-native, misfolded proteins and act upon them to prevent aggregation or to mitigate their toxic consequences (Hartl *et al*, 2011). This means that misfolded proteins are usually refolded to their functional conformations or degraded by quality control mechanisms. When misfolded proteins evade these quality control systems, they form aggregates that are implicated in a number of protein misfolding diseases (Hipp *et al*, 2014). More recently, the organized sequestration of misfolded proteins to defined inclusion sites has been recognized as a regulated process that depends on dedicated molecular chaperones, termed sequestrases (Hill *et al*, 2017; Miller *et al*, 2015; Mogk & Bukau, 2017). The sequestration of misfolded proteins into intracellular deposit sites helps cells to cope with an accumulation of misfolded proteins by partitioning them away from their normal productive pathways, protecting against potential cytotoxic effects and by facilitating targeted degradation (Gallardo *et al*, 2021). Although aggregation is a well-studied phenomenon and many key players are known, it remains unclear exactly how different growth and stress conditions cause protein aggregation and the degree of stress specificity in the chaperone response to aggregate formation is unknown.

The spatial sequestration of misfolded proteins is a highly conserved protein quality control (PQC) strategy, but the subcellular localization of deposition sites differs between organisms (Iburg *et al*, 2020; Kaganovich *et al*, 2008; Ogrodnik *et al*, 2014). The yeast model is currently the best characterized system relating to stress-induced protein aggregation where extensive genetic and cell biological analyses exist. Proteotoxic stress causes the formation of spatially separated protein deposits including IPOD (Insoluble Protein Deposit), CytoQ (cytosolic quality control compartment, Q bodies) and INQ (intranuclear quality control compartment, JUNQ) (Escusa-Toret *et al*, 2013; Kaganovich *et al*., 2008; Mogk & Bukau, 2017; Rothe *et al*, 2018; Sontag *et al*, 2017; Specht *et al*, 2011). Following protein misfolding, multiple CyoQs are formed in the cytoplasm, which are resolved into other deposition sites including INQ and IPOD (Kaganovich *et al*., 2008; Miller *et al*., 2015; Rothe *et al*., 2018). Cytosolic IPOD, CytoQ and nuclear INQ are thought to represent independent aggregate deposits which protect against overloading the proteostasis machinery during protein misfolding conditions (Miller *et al*., 2015). Proteins targeted to CytoQ and INQ can undergo refolding, whereas terminally misfolded and amyloid aggregates are thought to be targeted to the IPOD (Kaganovich *et al*., 2008; Miller *et al*., 2015)

Hsp42 and Btn2 are two key sequestrases required for the deposition of misfolded proteins into CytoQ and INQ, respectively (Malinovska *et al*, 2012; Miller *et al*., 2015; Specht *et al*., 2011). The cytosolic Hsp42 is a member of the small heat-shock family (sHSP) of chaperones and contains a prion-like domain (PrLD) that is essential for its aggregase function (Grousl *et al*, 2018). Like other members of the conserved sHSP family, Hsp42 forms a large homo-oligomeric structure and contains a highly conserved alpha-crystallin domain. Mutations in human sHSPs have been linked to various cardiovascular and neuromuscular diseases and hereditary cataracts. Hsp42 has been shown to direct protein sequestration to CytoQ during heat stress conditions using model fluorescently misfolded reporters (Grousl *et al*., 2018; Malinovska *et al*., 2012; Miller *et al*., 2015; Specht *et al*., 2011). The nuclear equivalent Btn2 is a small heat shock-like protein that is essential for INQ formation. Its protein levels are strongly induced by heat stress, and it functions in recruiting Hsp70/Hsp100 disaggregases for refolding of sequestered proteins during stress recovery (Ho *et al*, 2019; Malinovska *et al*., 2012; Miller *et al*., 2015). Btn2 has been shown to form high-molecular-weight complexes reminiscent of sHsp oligomers and similarly exhibits chaperone activity by associating with misfolded reporter proteins to increase their reactivation by Hsp70-Hsp100 chaperones (Ho *et al*., 2019). Yeast Btn2 has similarity to Hook1, a coiled-coil protein that associates with the cytoskeleton in mammalian cells. Its name derives from the finding that it is upregulated in response to loss of *BTN1*, which encodes an ortholog of a human Batten disease protein implicated in progressive neurodegeneration and early death.

Despite the established roles for Hsp42 and Btn2 as cellular sequestrases, their exact intracellular functions have remained elusive, especially since no growth defects have been observed in deletion mutants including a lack of sensitivity to heat stress (Haslbeck *et al*, 2004; Ho *et al*., 2019; Miller *et al*., 2015). In this current study we have investigated the roles of the Hsp42 and Btn2 protein sequestrases in maintaining proteostasis during oxidative stress conditions. Oxidative stress induced by hydrogen peroxide exposure is known to inhibit translation whilst increasing protein misfolding and aggregation (Shenton & Grant, 2003; Weids *et al*, 2016). We show that the levels of protein oxidative damage formed in response to oxidative stress are similar in wild-type and sequestrase mutants, but protein aggregation is elevated suggesting that the Btn2 and Hsp42 sequestrases normally act to sequester oxidatively damaged proteins as part of the cells antioxidant defence system. In agreement with this idea, we show that *btn2 hsp42* mutants are sensitive to hydrogen peroxide stress implicating a functional role for protein sequestration in oxidant tolerance. Together, our data show that the Btn2 and hsp42 sequestrases act to protect against widespread amorphous and amyloid protein aggregation during oxidative stress conditions.

## Results

### Btn2 and Hsp42 are required for tolerance to oxidative stress

Given the lack of sensitivity of sequestrase mutants to heat stress conditions, we examined whether Btn2 and Hsp42 are required for tolerance to oxidative stress, as another stress condition that causes protein misfolding and aggregation. For these experiments we used single *hsp42* and *btn2* mutants as well as a double *btn2 hsp42* mutant. We first confirmed that mutants lacking *BTN2* or *HSP42* are unaffected in temperature sensitivity (Fig. 1A). Interestingly, the *btn2* and *hsp42* mutants were sensitive to hydrogen peroxide and the double *btn2 hsp42* mutant showed strong sensitivity implicating a functional requirement for these chaperones during oxidative stress conditions. We also found that *btn2* and *btn2 hsp42* double mutants grow poorly under respiratory growth conditions suggesting a sensitivity to endogenously generated ROS (Fig. 1A). Cur1 is the yeast paralog of Btn2 but is not required for INQ formation (Malinovska *et al*., 2012). We examined the oxidant sensitivity of *cur1* and *btn2 cur1* mutants but found that loss of *CUR1* does not affect oxidant tolerance (Fig. 1A). The rest of this manuscript therefore focusses on the requirement for Btn2 and Hsp42 during oxidative stress conditions.

**Fig. 1.**
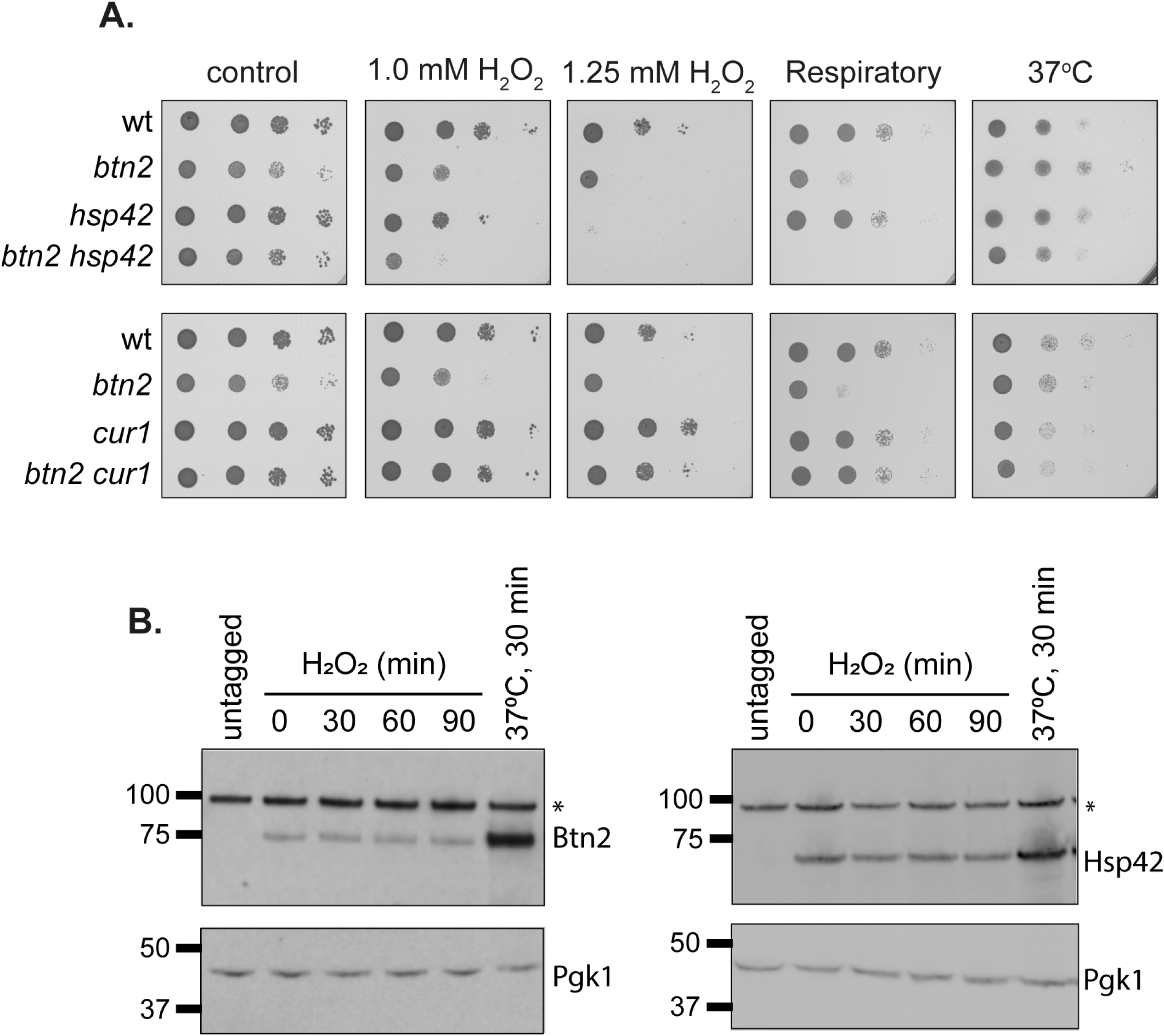
Mutants lacking Btn2 and Hsp42 and are sensitive to oxidative stress. **A.** The indicated strains were grown to exponential phase and the A_600_ adjusted to 1, 0.1, 0.01 or 0.001 before spotting onto plates. This included SD (control), hydrogen peroxide (1.0 and 1.25 mM), glycerol / ethanol (respiratory) and high temperature (37°C). **B.** Btn2 and Hsp42 protein levels are unaffected in response to oxidative stress conditions. Whole cell extracts were prepared from wild-type strains containing Btn2-Myc or Hsp42-Myc grown under non-stress conditions, subjected to a 37oC, 30-minute heat shock or exposed to 0.8 mM hydrogen peroxide for 30, 60 or 90 minutes. Western blots are shown probed with αMyc or α-Pgk1 as a loading control. Asterisks denote a non-specific band recognized by αMyc.

Despite not being required for heat tolerance, Btn2 and Hsp42 protein levels are increased in response to heat stress (Malinovska *et al*., 2012; Miller *et al*., 2015). Since we found that Btn2 and Hsp42 are required to promote oxidant tolerance, we tested whether their expression levels are also increased in response to oxidative stress conditions. However, no increases in Btn2 or Hsp42 protein levels were observed in response to hydrogen peroxide stress suggesting that the basal levels of these sequestrases are sufficient to promote oxidant tolerance (Fig. 1B). In comparison, Btn2 and Hsp42 expression levels were strongly induced in response to a 37^0^C heat stress (Fig. 1B).

### Increased protein oxidation does not account for the sensitivity of sequestrase mutants to oxidative stress

One possibility to explain the oxidant sensitivity of the *btn2 hsp42* double mutant is that protein oxidation is increased in sequestrase mutants. We examined protein carbonylation as a commonly used marker of protein oxidative damage (Nystrom, 2005). Carbonyl groups on proteins can be detected by immunoblot analysis using an antibody that recognizes the carbonyl-specific probe DNPH. Using this assay, we found that protein carbonylation is elevated by approximately 50% in a wild-type strain in response to oxidative stress caused by hydrogen peroxide exposure (Fig. 2A). The basal levels of protein carbonylation were increased in the *btn2, hsp42* and *btn2 hsp42* mutants during normal growth conditions. However, no further increase in protein oxidation was detected in sequestrase mutants compared with the wild-type strain during oxidative stress conditions (Fig. 2A). These data indicate that whilst protein oxidation is elevated in sequestrase mutants during non-stress conditions, elevated levels of oxidatively damaged proteins do not appear to accumulate in sequestrase mutants that might explain the sensitivity of the double *btn2 hsp42* mutant to oxidative stress.

**Fig. 2.**
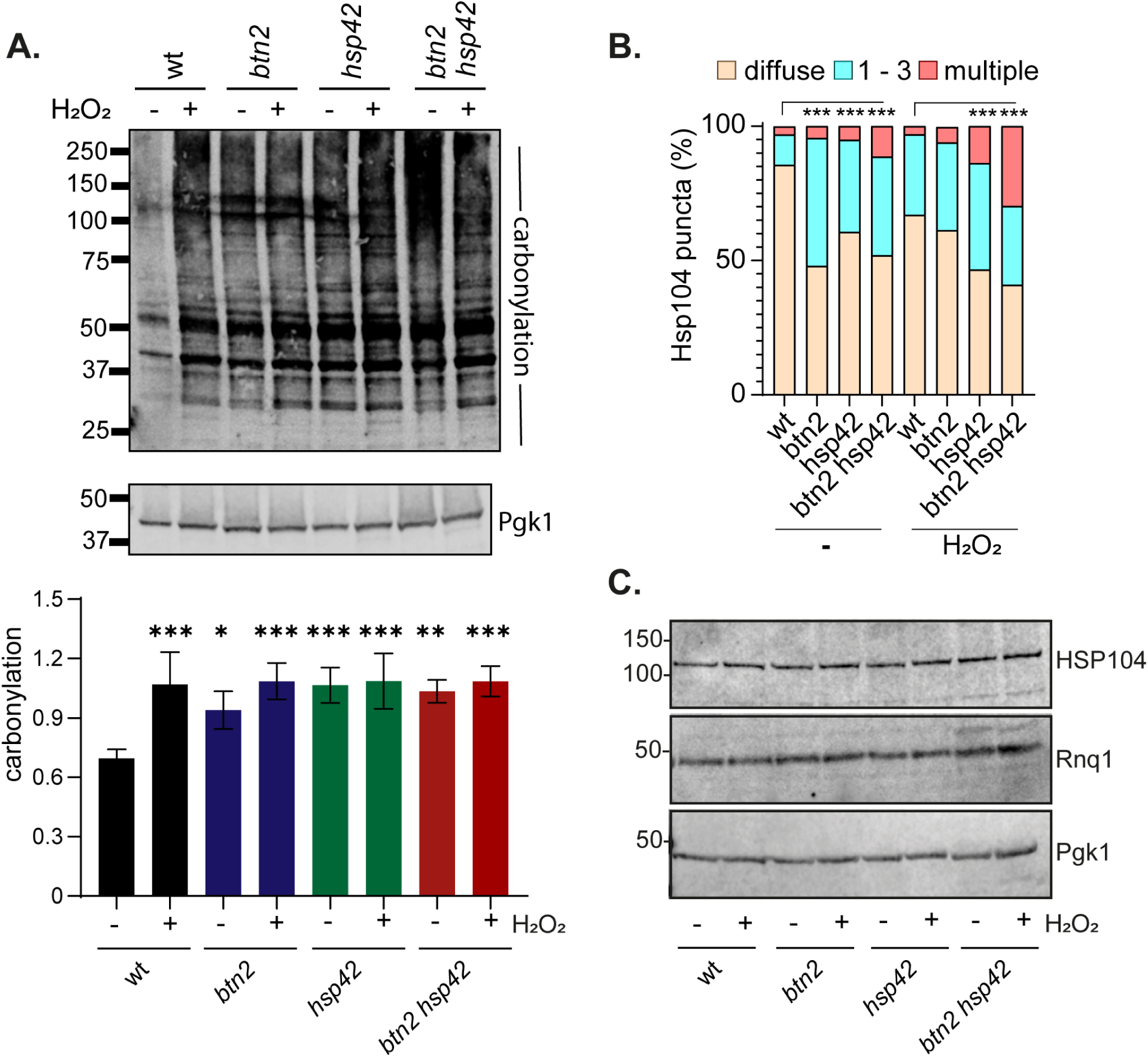
Protein aggregation is increased in sequestrase mutants. **A.** Wild-type and sequestrase mutants were grown to exponential phase and treated with 0.8 mM hydrogen peroxide (+) or left untreated (-) for one hour. Protein extracts were treated with the carbonyl-specific probe, DNPH, and analyzed by Western blot analysis using an antibody against DNPH. Quantitative data is shown as the means of three independent biological repeat experiments (carbonylation relative to Pgk1). All data are an average of n ≥ 3 replicates ± SD; * p<0.05, * p<0.01, * p<0.001 (one-way ANOVA). **B.** Hsp104-RFP was visualized in wild-type and sequestrase mutant cells grown to exponential phase and treated with 0.8 mM hydrogen peroxide or left untreated for one hour. Charts show the percentage of cells contain 0, 1-3, or >3 punct per cell scored in 300 cells for each strain. Significance is shown compared with the wild-type strain; *** *p* < 0.001 (Mann–Whiney U-test). **C.** Western blot analysis blot analysis of the same strains probed with antibodies that recognize Hsp104, Rnq1 or Pgk1.

### Oxidative stress-induced protein aggregation is increased in sequestrase mutants

We next examined whether protein aggregation is altered in sequestrase mutants to explain their sensitivity to oxidative stress. Hsp104 is the main cellular disaggregase that mediates refolding from the aggregated state and can be used to visualize the sites of protein aggregate formation in cells (Erjavec *et al*, 2007; Glover & Lindquist, 1998; Hamdan *et al*, 2017; Lee *et al*, 2010). We hypothesized that Hsp42 and Btn2-mediated sequestration might normally act to triage oxidized proteins and their absence would therefore lead to an accumulation of protein aggregates in *hsp42 btn2* mutants. Approximately 15% of wild-type cells were found to contain visible Hsp104-puncta during normal growth conditions (Fig. 2B). Protein aggregation was significantly increased in response to oxidative stress with greater than 30% of cells containing protein aggregates following exposure to hydrogen peroxide. The number of cells containing Hsp104-marked aggregates was increased in all sequestrase mutants (*btn2, hsp42, btn2 hsp42*) during non-stress conditions. For the *hsp42* and *btn2 hsp42* mutants, protein aggregation was further increased in response to oxidative stress compared with the wild-type during oxidative stress conditions (Fig. 2B). This was particularly apparent for the *btn2 hsp42* mutant strains where 30% cells contained multiple Hsp104-marked aggregates.

Given the increase in Hsp104-marked aggregates, we examined the cellular concentrations of Hsp104 to determine whether the increased puncta formation observed in sequestrase mutants correlates with increased expression of Hsp104 (Fig. 2C). However, the cellular concentrations of Hsp104 were comparable in all strains during both non-stress and oxidative stress conditions suggesting that the increased Hsp104 puncta formation arises due to re-localization of existing pools of Hsp104 in sequestrase mutants rather than through new chaperone synthesis.

### Btn2 and hsp42 co-localize with Hsp104-marked protein aggregates following oxidative stress conditions

Hsp104 has been variously localized to INQ, CytoQ and IPOD following heat-induced-protein misfolding (Ho *et al*., 2019; Kaganovich *et al*., 2008; Lee *et al*, 2018; Malinovska *et al*., 2012; Specht *et al*., 2011). Given the increase in Hsp104-marked protein aggregates following oxidative stress, we examined whether the co-localization of Hsp104 with Btn2 or Hsp42 is affected during these conditions. If Btn2 and Hsp42 are required to triage oxidatively damaged proteins, we reasoned that Hsp104-marked aggregates would co-localize with these sequestrases. For these experiments we expressed Hsp104-RFP in wild-type strains that express Btn2-GFP or Hsp42-GFP. Co-localization of both Hsp42 and Btn2 was observed with Hsp104 during normal non-stress conditions, and this was significantly increased following hydrogen peroxide exposure for both sequestrases suggesting that Btn2 and Hsp42 play a role in triaging aggregated proteins formed during oxidative stress conditions (Fig. 3A).

**Fig. 3.**
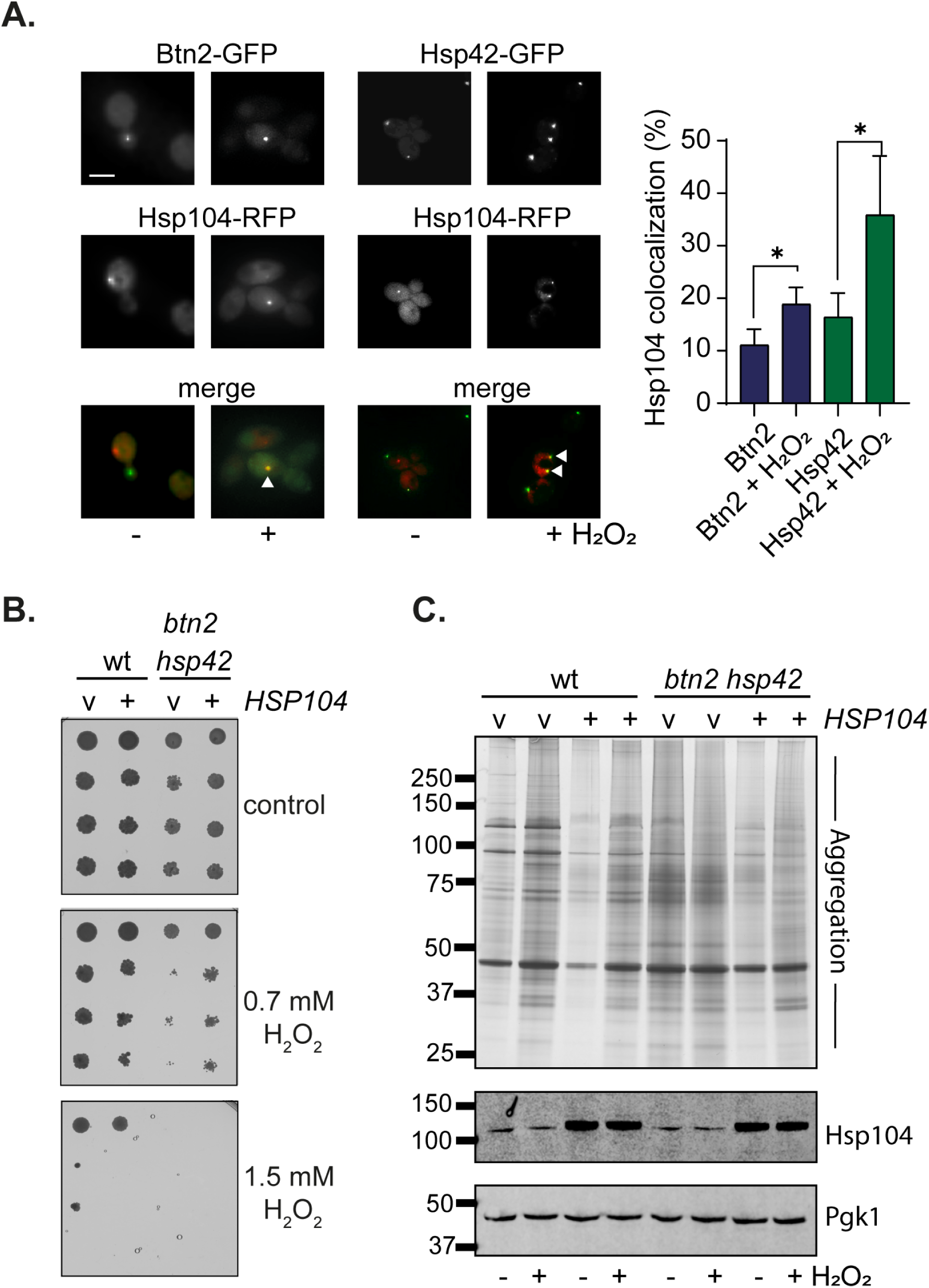
Hsp104 is required for oxidative stress tolerance in sequestrase mutants. **A.** Representative epifluorescent microscopic images are shown from strains expressing Btn2-GFP or Hsp42-GFP and Hsp104-RFP. Strains were grown to exponential phase and left untreated or treated with with 0.8 mM hydrogen peroxide for one hour. Quantification is for the co-localisation (%) of Btn2 or Hsp42 puncta with Hsp104 puncta from three biological replicates. Error bars denote SD and significance is shown compared with the untreated strains, * *p* < 0.05 (t-test). **B.** Overexpression of Hsp104 improves the hydrogen peroxide sensitivity of *btn2 hsp42* mutants. The wild-type and *btn2 hsp42* mutant strains containing vector control or expressing *HSP104* under the control of the constitutive *TDH3* promoter were grown to exponential phase and the A_600_ adjusted to 1, 0.1, 0.01 or 0.001 before spotting onto SD plates containing the indicated concentrations of hydrogen peroxide. **C.** Protein aggregates were isolated from the wild-type and *btn2 hsp42* mutant strains containing vector control or overexpressing *HSP104* and analysed by SDS-PAGE and silver staining. Western blot analysis of the same strains probed with α-Hsp104 or α-Pgk1.

### Overexpression of Hsp104 rescues the oxidant sensitivity of sequestrase mutants

Since Hsp104 appears to be required to protect against protein aggregation during the response to oxidative stress, but its cellular concentrations remain unaltered during these stress conditions, we hypothesised that Hsp104 may become limiting in *btn2 hsp42* mutants. To test this idea, *HSP104 was* over-expressed under the control of the constitutively active TDH3 promoter (Malinovska *et al*., 2012) in wild-type and *btn2 hsp42* mutant strains. Whilst overexpression of *HSP104* did not improve the hydrogen peroxide tolerance of the wild-type strain, the hydrogen peroxide sensitivity of the *btn2 hsp42* mutant was improved (Fig. 3B). This suggests that Hsp104 can become limiting in sequestrase mutant during oxidative stress conditions that promote protein misfolding and aggregation.

To further confirm that overexpression of Hsp104 rescues the oxidant sensitivity of *btn2 hsp42* mutants by protecting against the toxicity of protein aggregate accumulation in the absence of sequestrase activity, we directly examined protein aggregation in wild-type and sequestrase mutants. Insoluble protein aggregates were separated from soluble proteins by differential centrifugation and any contaminating membrane proteins removed by detergent washes (Hamdan *et al*., 2017; Jang *et al*, 2004; Weids & Grant, 2014). Low levels of protein aggregation were detected in the wild-type strain during non-stress conditions which were increased in response to hydrogen peroxide exposure (Fig. 3C). Protein aggregation was decreased by overexpression of Hsp104 under both non-stress and oxidative stress conditions consistent with the function of hsp104 as a disaggregase. Protein aggregation was strongly increased in the *btn2 hsp42* mutant and again Hsp104 overexpression reduced the cellular levels of protein aggregation. Hsp104 overexpression also appeared to cause alterations in the protein aggregates detected in the *btn2 hsp42* mutant particularly under oxidative stress (Fig. 3C). Western blotting was used to confirm that Hsp104 is similarly overexpressed in wild-type and *btn2 hsp42* mutants (Fig. 3C). Taken together, these data indicate that Btn2 and Hsp42 are required to protect against protein aggregation caused by oxidative stress conditions in a mechanism that requires the Hsp104 disaggregase.

### Sup35 localizes with Btn2 and Hsp42 during non-stress and oxidative stress conditions

Most studies that have examined protein misfolding and aggregation in sequestrase mutants have relied on model PQC reporter substrates and little is known regarding the *in vivo* substrates of these chaperones. We decided to examine aggregation of the Sup35 eukaryotic release factor 3 (eRF3) in sequestrase mutants as a potential chaperone substrate. This is because hydrogen peroxide exposure has been shown to cause extensive aggregation of Sup35 in cells (Jamar *et al*, 2017, 2018). Sup35 is also well known for its ability to form prion aggregates known as [*PSI^+^*] and is commonly used as a model to study amyloidogenic aggregation (Wickner, 1994). Importantly, oxidative damage to the non-prion form of Sup35 has been shown to be an important trigger influencing the formation of heritable [*PSI*^+^] prions in cells (Doronina *et al*, 2015; Grant, 2015; Sideri *et al*, 2011; Sideri *et al*, 2010; Tyedmers *et al*, 2008).

Previous studies have described a high level of Btn2 colocalization with Sup35 aggregates formed in [*PSI*^+^] strains (Kryndushkin *et al*, 2008). Additionally, overexpression of *NM-SUP35-GFP* (a fusion between the N-terminal prion domain (PrD) and the middle M domain of Sup35 with GFP) has been shown to result in the formation of multiple Sup35-marked puncta where a single puncta commonly co-localizes with Hsp42 (Arslan *et al*, 2015). We wanted to test whether Sup35 localizes with the Btn2 or Hsp42 sequestrases during oxidative stress conditions. For these experiments, we transiently expressed Sup35-NM-RFP for two hours under the control of the inducible *GAL1* promoter in wild-type [*psi*^-^] cells. Coalescence of newly made Sup35-NM-RFP with pre-existing Sup35 aggregates allows the detection of Sup35 protein aggregates in cells (Patino *et al*, 1995). During non-stress conditions, approximately 40% of cells contained visible Sup35 puncta, and the number of puncta was increased in response to oxidative stress (Fig. 4A and B). Similarly, the number of cells containing Hsp42 or Btn2-marked puncta was also found to increase in response to hydrogen peroxide exposure (Fig. 4A and B). Btn2 and Hsp42 colocalized with approximately 35% and 15% of Sup35 puncta, respectively, during both non-stress and oxidative stress conditions (Fig. 4C). This suggests that the extent of colocalization of Sup35 with Btn2 and Hsp42 is maintained during oxidative stress conditions along with the increase in Sup35, Btn2 and Hsp42 puncta.

**Fig. 4.**
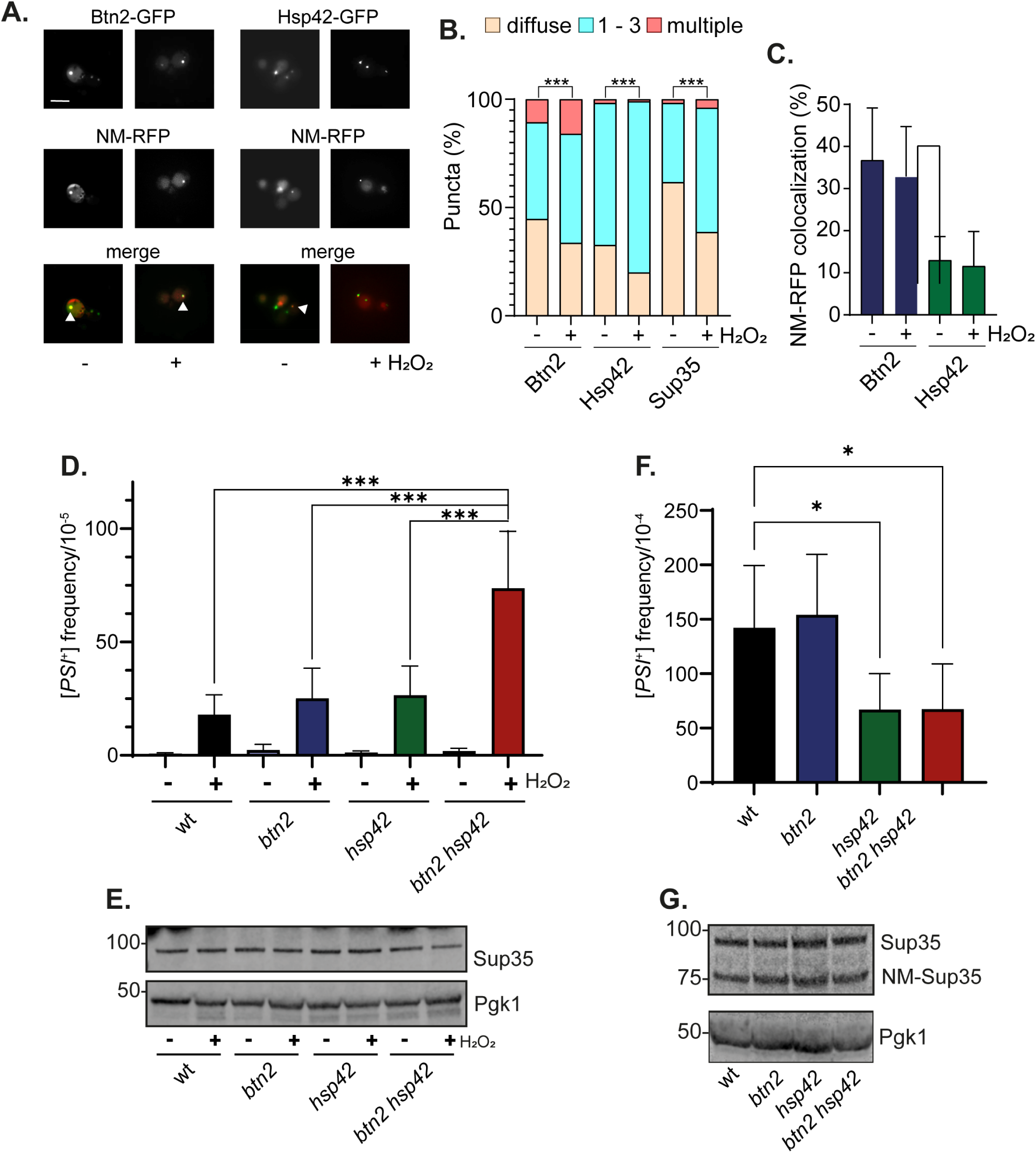
Sup35 localizes with Btn2 and Hsp42 and oxidant-induced [*PSI*^+^] prion formation is increased in mutants lacking *BTN2* and *HSP42.* **A.** Representative epifluorescent microscopic images are shown from strains expressing Btn2-GFP or Hsp42-GFP and NM-RFP. Strains were grown to exponential phase and left untreated or treated with with 0.8 mM hydrogen peroxide for one hour. Sup35-NM-RFP was induced for two hours under the control of the inducible *GAL1* promoter. White arrows indicate examples of colocalization. **B.** Charts show the percentage of cells that contain 0, 1-3, or >3 Btn2, Hsp42 or Sup35 puncta per cell scored in 300 cells for each strain. Significance is shown comparing stressed and unstressed strains; *** *p* < 0.001 (Mann–Whiney U-test). **C.** Quantification is shown for the co-localisation of Btn2 or Hsp42 (%) with Sup35 puncta from three biological replicates. Error bars denote SD. **D.** [*PSI*^+^] prion formation was quantified in the wild-type, *btn2, hsp42* and *btn2 hsp42* mutant strains during non-stress and oxidative stress conditions. Data shown are the means of at least three independent biological repeat experiments expressed as the number of colonies per 10^5^ viable cells. Error bars denote standard deviation. Significance is shown using a one-way ANOVA test; *** *p* < 0.001. **E.** Western blot analysis of the same strains as for A. probed with α-Sup35 or α-Pgk1 as a loading control. **F.** [*PSI*^+^] prion formation was quantified in the wild-type and indicated mutant strains containing the *Sup35NM-GFP* plasmid following 20 hours of copper induction. Data shown are the means of at least three independent biological repeat experiments expressed as the number of colonies per 10^4^ viable cells. Error bars denote standard deviation; * marks statistical significance at p<0.01 (one-way ANOVA). **G.** Western blot analysis of the same strains as for C. probed with α-Sup35 or α-Pgk1.

### Oxidant-induced prion formation is increased in *btn2 hsp42* sequestrase mutants

We next examined [*PSI*^+^] prion formation as a measure of amyloidogenic aggregation in sequestrase mutants. We found that the frequency of [*PSI*^+^] prion formation is unaffected in *btn2* or *hsp42* single and double mutants during normal non-stress growth conditions indicating that the Btn2 and hsp42 sequestrases do not play an anti-prion role in in suppressing prion formation during non-stress conditions (Fig. 4D). Following oxidative stress, the frequency of [*PSI*^+^] formation was increased by greater than 20-fold and similar increases were observed in the *btn2* and *hsp42* single mutants. Interestingly however, the frequency of oxidant-induced [*PSI*^+^] formation was significantly increased in the *btn2 hsp42* double mutant compared with the wild-type and single mutant strains (Fig. 4D). Given that increased cellular concentrations of Sup35 can promote [*PSI*^+^] prion formation, the levels of Sup35 were measured in all mutant strains (Fig. 4E). This analysis confirmed that similar levels of Sup35 protein are present ruling out any effects on Sup35 protein concentrations.

The best-established method to increase the frequency of *de novo* [*PSI*^+^] formation is to overexpress Sup35 which increases the probability of soluble Sup35 switching to its prion form (Wickner, 1994). We therefore examined whether loss or *BTN2* or *HSP42* influences overexpression-induced [*PSI*^+^] prion formation. The frequency of [*PSI*^+^] prion formation was strongly induced in the wild-type strain following overexpression of *NM-SUP35-GFP* for 20 hours as expected (Fig. 4F). However, no significant differences were observed in the *btn2* mutant, and overexpression induced [*PSI*^+^] prion formation was somewhat reduced in the *hsp42* and *btn2 hsp42* mutants. Immunoblotting was used to confirm that similar Sup35 and NM-GFP protein concentrations are present in all strains (Fig. 4G). Taken together, these data indicate that Btn2 and Hsp42 suppress the frequency of oxidant-induced, but not overexpression*-*induced, [*PSI*^+^] prion formation.

### Sup35 aggregate formation is increased in sequestrase mutants

To address how the loss of both Btn2 and Hsp42 might influence oxidant induced [*PSI^+^*] formation, we examined Sup35 and Rnq1 aggregation in sequestrase mutants. *De novo* formation of the [*PSI^+^*] prion requires the presence of a second prion called [*PIN^+^*] for [*PSI*^+^] inducibility, which is usually the prion form of Rnq1, a protein of unknown function (Derkatch *et al*, 2001; Derkatch *et al*, 1997). [*PIN^+^*] prion aggregates can cross-seed [*PSI*^+^] formation by acting as templates on which Sup35 molecules misfold and assemble into prion aggregates (Derkatch *et al*., 2001; Derkatch *et al*, 2004). Rnq1 localizes to the IPOD which has been proposed to be the site where irreversibly aggregated proteins are triaged and to act as a site for *de novo* prion formation (Tyedmers *et al*, 2010). Accordingly, colocalization of Sup35 with Rnq1 at the IPOD has been observed during both overexpression and oxidative stress induced [*PSI*^+^] prion formation (Arslan *et al*., 2015; Speldewinde *et al*, 2017).

To examine whether Sup35 or Rnq1 aggregate formation and localization is affected in mutants lacking *HSP42* and *BTN2,* we expressed *RNQ1-CFP* in wild-type and sequestrase mutants expressing *SUP35-GFP* as the sole copy of *SUP35* under the control of its endogenous promoter. We found that most wild-type cells with visible Sup35-GFP marked puncta contained 1-3 puncta per cell, and the number of cells containing Sup35-GFP puncta was increased in response to oxidative stress (Fig. 55A and B). No significant differences in the pattern of Sup35-GFP puncta were observed in the *btn2* or *hsp42* mutants. In contrast, Sup35-GFP puncta formation was significantly elevated in the *btn2 hsp42* mutant with more cells containing visible puncta which were often present as multiple fainter Sup35 puncta in cells and puncta formation was further increased in response to oxidative stress (Fig. 5A and B).

**Fig. 5.**
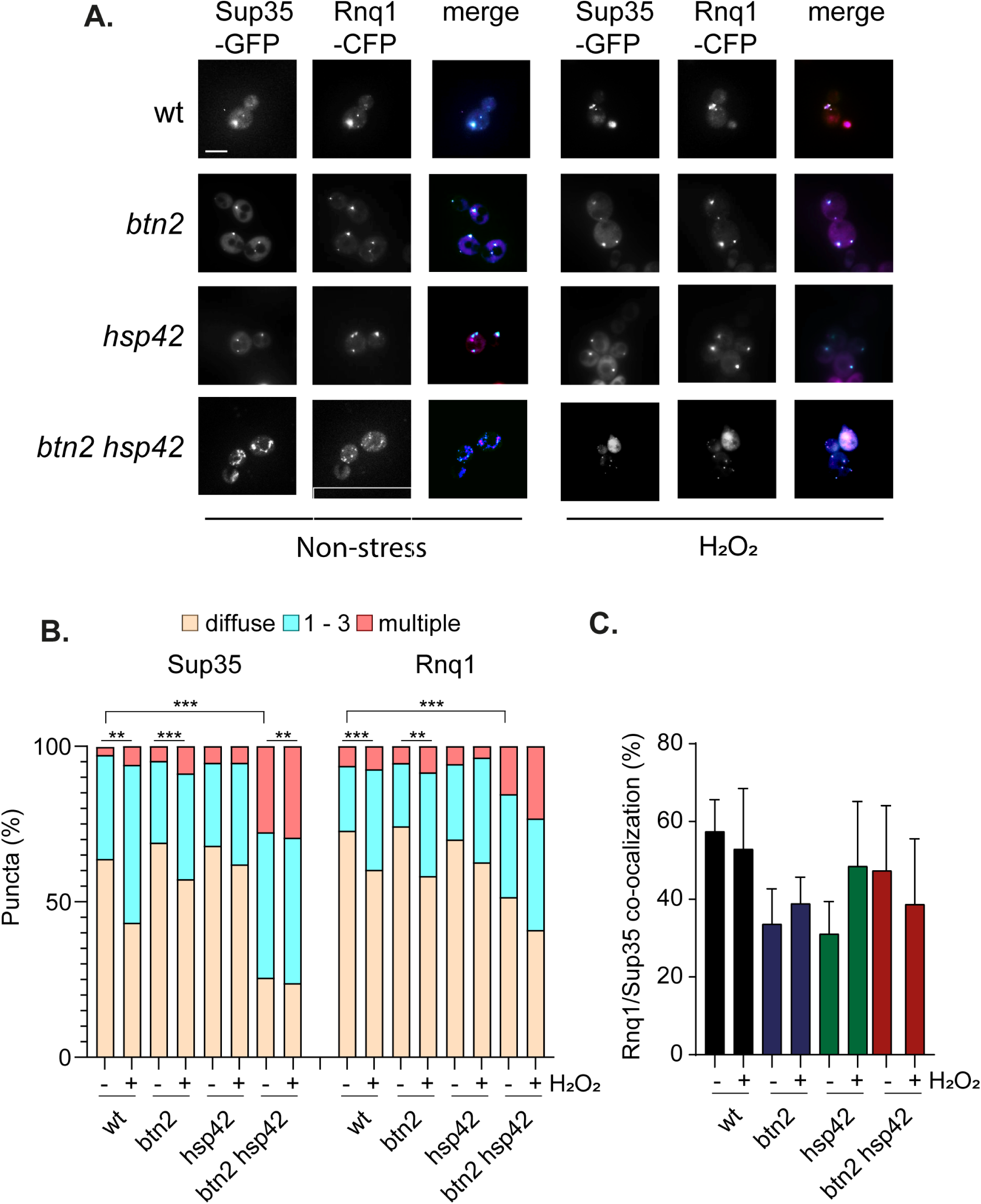
Sup35 aggregate formation is increased in sequestrase mutants. **A.** Representative epifluorescent microscopic images are shown from strains expressing Sup35-GFP and Rnq1-CFP. Strains were grown to exponential phase and left untreated (non-stress) or treated with with 0.8 mM hydrogen peroxide for one hour. GFP is false coloured magenta and CFP is false coloured cyan. **B.** Charts show the percentage of cells contain 0, 1-3, or >3 Sup35 or Rnq1 puncta per cell scored in 300 cells for each strain. Significance: ** *p* < 0.01, *** *p* < 0.001 (Mann–Whiney U-test). **C.** Quantification is shown for the co-localisation of Rnq1 with Sup35 puncta from three biological replicates. Error bars denote SD.

Distinct patterns of Rnq1-GFP fluorescence have been described in [*PIN*^+^] cells which have been referred to as single-dot (s.d.) or multiple-dot (m.d.) depending on the numbers of fluorescent puncta detected (Bradley & Liebman, 2003). Multiple dots have also been observed which are smaller and fainter in intensity (Huang *et al*, 2013). This pattern of Rnq1 puncta tends to correlate with strong [*PIN*^+^] variants suggesting that multiple dots are associated with more heritable prion seeds (Bradley *et al*, 2002; Bradley & Liebman, 2003; Huang *et al*., 2013). Most wild-type cells with visible Rnq1-CFP marked puncta contained 1-3 puncta per cell, which was increased in response to oxidative stress (Fig. 5A). Like Sup35, no significant differences in the pattern of Rnq1 puncta were observed in the *btn2* or *hsp42* single mutants. However, Rnq1 puncta formation was also increased in the *btn2 hsp42* mutant with more cells containing visible puncta which were often present as multiple fainter puncta in cells (Fig. 5A). We used immunoblotting to confirm that there are no differences in Rnq1 protein levels in sequestrase mutants that might account for the altered pattern of Rnq1 puncta formation observed using microscopy (Fig. 2C).

When we examined the co-localization of Rnq1 with Sup35, we found that Rnq1-CFP puncta co-localized with approximately 60% of Sup35 puncta in the wild-type strain and this was unaffected by oxidative stress (Fig. 5C). Similar levels of Rnq1 co-localization with Sup35 were detected in the *btn2, hsp42* or *btn2 hsp42* mutant under non-stress and oxidative stress conditions. Taken together, these data indicate that the number of cells containing Sup35 and Rnq1 aggregates is increased in the *btn2 hsp42* mutant compared with the wild-type strain and a further increase in Sup35 aggregation is observed in the *btn2 hsp42* mutant in response to oxidative stress correlating with the increased prion formation observed in this mutant. Despite the alterations in Sup35 and Rnq1 puncta formation, the extent of Sup35-Rnq1 colocalization is maintained at a consistent level.

### IPOD localization is not required for increased oxidant-induced prion formation in *btn2* hsp42 mutants

The IPOD is thought to promote prion formation by acting as a site where the localised concentration of misfolded proteins acts to facilitate the nucleation of prion protein polymerisation (Tyedmers *et al*., 2010). The amyloidogenic [*PIN*^+^] prion form of Rnq1 localizes to the IPOD and is often used to visualize the IPOD which is usually present as a single large perivacuolar inclusion site in cells (Sontag *et al*, 2014; Tyedmers *et al*., 2010). Hence, our finding that multiple small aggregation sites containing Rnq1 and Sup35 are formed in *btn2 hsp42* mutants suggests that IPOD localization may not be required for the increased [*PSI*+] formation observed in this mutant. To test the requirement for IPOD localization, we quantified the frequency of [*PSI*+] formation in *btn2 hsp42* mutants lacking *ABP1.* Loss of *ABP1* has been shown to disrupt the cortical actin cytoskeleton which is required to mediate the IPOD localization of oxidized Sup35 (Speldewinde *et al*., 2017). We first confirmed that the increased frequency of [*PSI*^+^] formation induced in response to hydrogen peroxide treatment is abrogated in an *abp1* mutant (Fig.6A). In contrast, loss of *ABP1* did not affect the frequency of oxidant-induced [*PSI*^+^] formation in *btn2 hsp42* mutants confirming that IPOD localization is not required for [*PSI*^+^] formation in this mutant (Fig. 6A). This indicates that prion formation is likely occurring at multiple other sites in *btn2 hsp42* mutants during oxidative stress conditions.

**Fig. 6.**
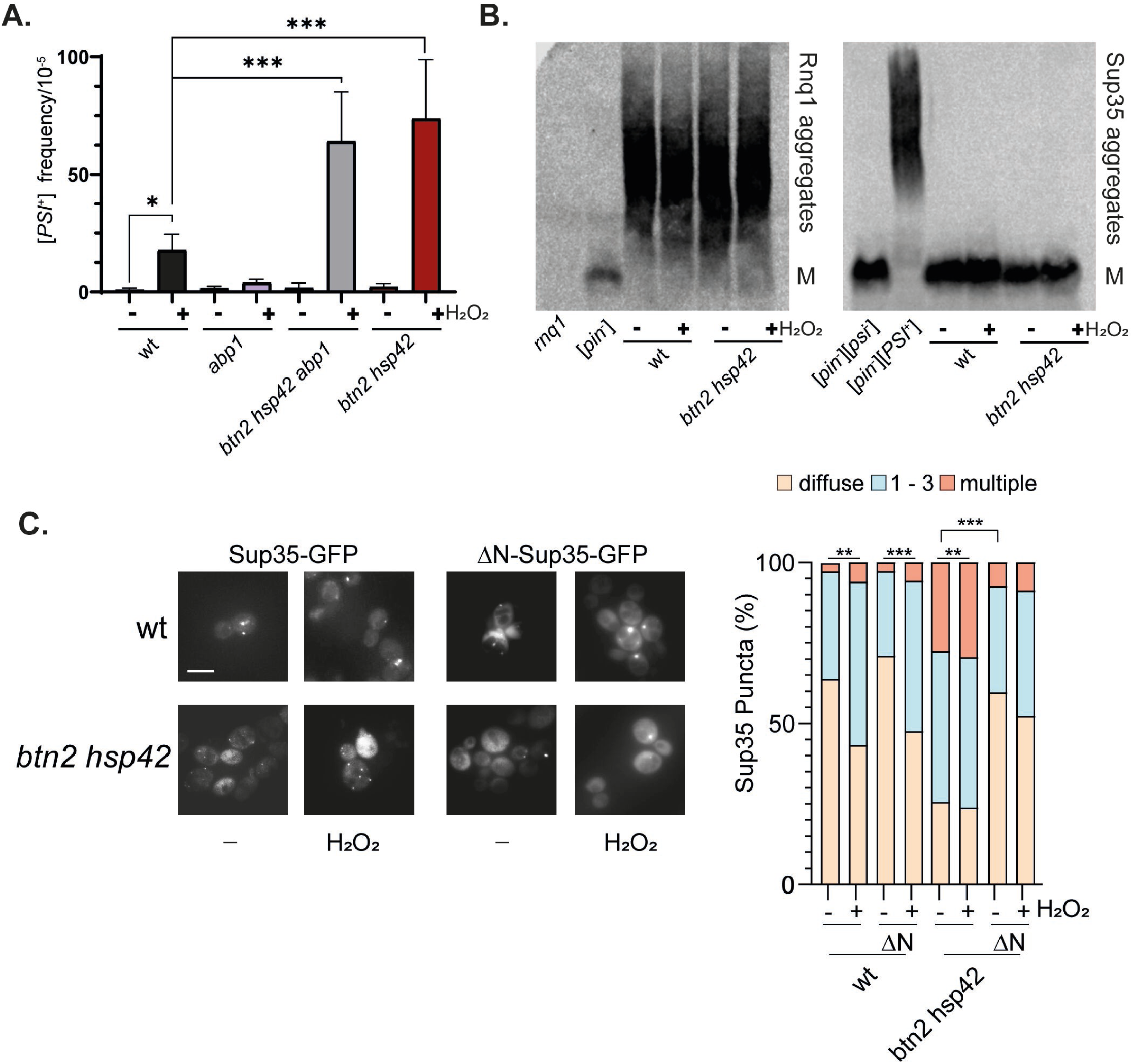
Non-amyloid aggregate formation underlies the increased Sup35 aggregation observed in *btn2* hsp42 mutants. **A.** [*PSI*^+^] prion formation was quantified in the wild-type, *abp1, btn2 hsp42* and *btn2 hsp42 abp1* mutant strains during non-stress conditions and following exposure to 0.8 mM hydrogen peroxide for one hour. Data shown are the means of at least three independent biological repeat experiments expressed as the number of colonies per 10^5^ viable cells. Error bars denote standard deviation. Significance is shown using a one-way ANOVA test; * *p* < 0.05 *** *p* < 0.001. **B.** SDS-resistant Sup35 and Rnq1 aggregates were detected in the wild-type and *btn2 hsp42* mutant strains using SDD-AGE. Strains were grown to exponential phase and left untreated (-) or treated with with 0.8 mM hydrogen peroxide (+) for one hour. [*psi*^-^] and *rnq1* deletion strains are shown for comparison with Rnq1 and [*PSI*^+^] and [*psi*^-^] strains are shown for comparison with Sup35. Aggregate and monomer (M) forms are indicated. **C.** Representative epifluorescent microscopic images are shown from strains expressing Sup35-GFP or ΔN-Sup35-GFP. Strains were grown to exponential phase and left untreated (non-stress) or treated with with 0.8 mM hydrogen peroxide for one hour. Charts show the percentage of cells contain 0, 1-3, or >3 Sup35 puncta per cell scored in 300 cells for each strain. Significance: ** *p* < 0.01, *** *p* < 0.001 (Mann–Whiney U-test).

### Non-amyloid aggregate formation underlies the increased Sup35 aggregation observed in *btn2* hsp42 mutants

Whilst the increased Sup35 puncta formation detected in *btn2 hsp42* mutants in response to oxidative stress appears to explain the increased [*PSI+*] formation, the number of fluorescent Sup35-GFP puncta observed is much higher than the frequency of [*PSI*^+^] prion formation. For example, whilst 75% of *btn2 hsp42* mutant cells contained at least one Sup35 puncta following oxidative stress conditions (Fig. 5B), the frequency of prion formation is approximately 8 x 10^-4^ under these conditions (Fig. 6A). This is generally observed since some cells containing aggregates may not be viable and many aggregates may not be amyloid aggregates (Arslan *et al*., 2015). We reasoned that the elevated puncta formation observed in the *btn2 hsp42* mutant may therefore reflect an increase in both amyloid and amorphous Sup35 aggregation. We therefore examined Sup35 and Rnq1 amyloid aggregation used semi-denaturing detergent-agarose gel electrophoresis (SDD-AGE) which can be used to separate monomeric Rnq1 or Sup35 from their high molecular weight SDS-resistant aggregate forms, diagnostic of amyloid formation (Bagriantsev & Liebman, 2004; Huang *et al*., 2013; Kryndushkin *et al*, 2003). SDS-AGE can also reveal differences in prion variants which display differences in size with weaker [*PSI*^+^] or [*PIN*^+^] variants forming larger protein aggregates compared with the smaller aggregates present in strong variants.

The *btn2 hsp42* mutant used in this study was constructed in a [*PIN*^+^][*psi*^-^] strain and we found that loss of both *BTN2* and *HSP42* did not affect Rnq1 aggregate sizes in the presence or absence of oxidative stress compared with the wild-type strain (Fig. 6B). This suggests that the multiple smaller Rnq1 puncta observed in the *btn2 hsp42* mutant do not arise due to altered amyloid formation. Whilst Sup35 amyloid aggregate formation was readily detectable in a control [*PSI*^+^] strain, no SDS-resistant aggregates were detected in the wild-type or *btn2 hsp42* mutant (Fig. 6B). These data suggest that the differences in Sup35 aggregate number and intensity observed in the *btn2 hsp42* mutant using microscopy do not predominantly arise due to increased amyloid formation and are most likely due to increased Sup35 amorphous aggregation.

To further investigate the nature of the aggregates present in the *btn2 hsp42* mutant, we examined Sup35 aggregation using a mutant lacking its N-terminal PrD. The PrD mutant of Sup35 (*ΔN-SUP35-GFP*) is sufficient to maintain viability but is deficient in prion propagation (Vishveshwara *et al*, 2009). The N-domain has also been shown to promote Sup35 phase separation and gelation forming non-amyloid aggregates during nutrient depletion stress conditions (Franzmann *et al*, 2018). When we expressed *ΔN-SUP35-GFP* as the sole copy of *SUP35* under the control of its endogenous promoter, we found that the number of Sup35 puncta in the wild-type strain was comparable for both wild-type Sup35 and Sup35 lacking its PrD indicating that the prion domain is not required for Sup35 puncta formation in wild-type strains (Fig. 6C). In contrast, loss of the PrD significantly decreased Sup35 puncta formation in the *btn2 hsp42* mutant indicating that the increased puncta formation in this mutant requires the PrD. This suggests that the increased Sup35 puncta formed in the *btn2 hsp42* mutant are reminiscent of the phase separated Sup35 condensates that are formed in response to energy depletion rather that amyloid-like prion particles (Franzmann *et al*., 2018).

## DISCUSSION

Many links have been established between protein aggregation and oxidative stress (Bossy-Wetzel *et al*, 2008; Levy *et al*, 2019). ROS have frequently been implicated in protein oxidative damage and partially misfolded proteins are known to be more susceptible to oxidation and aggregation (Dean *et al*, 1997). Newly synthesized proteins are particularly vulnerable to misfolding events and widespread aggregation is thought to be toxic, especially when the proteostasis network is compromised (Weids & Grant, 2014; Winkler *et al*, 2010). Protein aggregation is often accompanied by an increase in oxidative damage to cells and in many aggregation diseases, oxidative stress is an integral part of the pathology (Hands *et al*, 2011). Not surprisingly therefore, enzymes with antioxidant activity have been extensively linked with protein aggregation (Jang *et al*., 2004; Ling *et al*, 2012; MacDiarmid *et al*, 2013; Rand & Grant, 2006). Despite these established links between oxidative stress and protein aggregation, surprisingly little is known regarding the requirement for chaperones to mitigate the toxic effects of oxidative stress. Our data identify the Btn2 and Hsp42 sequestrases as key chaperones required for oxidant tolerance that are required to sequester misfolded proteins into defined PQC sites following ROS exposure.

The proper sequestration of aggregates is an important defence strategy that prevents the dysfunction and toxicity that is associated with protein misfolding diseases. Hsp42 has been shown to direct protein sequestration to CytoQ during heat stress conditions, whereas Btn2 is a small heat shock-like protein that is essential for INQ formation (Grousl *et al*., 2018; Ho *et al*., 2019; Malinovska *et al*., 2012; Miller *et al*., 2015; Specht *et al*., 2011). Despite the established roles for Hsp42 and Btn2 in protein sequestration during heat stress, their exact intracellular functions have remained elusive since they are not required for heat tolerance. Growth defects have been observed in mutants lacking the Btn2 and Hsp42 sequestrases by genetically limiting the capacity of Hsp70 (Ho *et al*., 2019). This suggests that sequestering unfolded proteins into defined deposit sites prevents titrating Hsp70 away from its essential chaperone functions in protein folding. The induction of Btn2 and Hsp42 expression in Hsp70 mutants may therefore act to counter limitations in Hsp70 capacity (Ho *et al*., 2019). The finding that mutants lacking Btn2 and Hsp42 are sensitive to hydrogen peroxide stress, but not heat stress, suggests a more direct functional requirement for these sequestrases during stress conditions that promote protein oxidation.

Hydrogen peroxide stress is known to inhibit translation whilst increasing protein aggregation (Shenton & Grant, 2003; Weids *et al*., 2016). All amino acids in proteins are potential targets of oxidation. They can be directly damaged on amino acid sidechains and backbone sites as well as through targeted oxidation of specific residues such as cysteine and methionine in proteins (Davies, 2016; Dean *et al*., 1997). Such modifications can significantly alter protein structure by affecting side-chain hydrophobicity, protein folding and amino acid interactions, often resulting in unfolding and protein aggregation. This contrasts with heat stress which is a well-characterized denaturing stress that causes protein unfolding and is generally reversible at non-extreme temperatures (Vabulas *et al*, 2010; Wallace *et al*, 2015). Hence, it is possible that the mechanistic differences in protein misfolding and aggregation caused by heat and oxidative stress accounts for the differential requirements for sequestrases during different stress conditions. The irreversible nature of many types of protein oxidation may also require sequestration as a strategy to prevent protein aggregates from accumulating at non-specific sites as we observed in *btn2 hsp42* mutants.

The sequestration of misfolded proteins is thought to protect the proteome against an overload of protein misfolding. Refolding from the aggregated state can be mediated by disaggregases including the AAA+ family Hsp104 together with Hsp70 and Hsp40 family members (Glover & Lindquist, 1998; Ho *et al*., 2019; Rampelt *et al*, 2012; Shorter, 2011). We found that Hsp104 co-localization with Btn2 and Hsp42 is increased in response to hydrogen peroxide exposure. However, Hsp104 localization, but not protein levels, was altered under these conditions suggesting that Hsp104 may become limiting in *btn2 hsp42* mutants in the face of widespread protein oxidation. In agreement with this idea, we found that overexpressing Hsp104 improved oxidant tolerance in *btn2 hsp42* mutants. Increasing Hsp104 levels in a wild-type strain did not affect oxidant tolerance indicating that Hsp104 levels are normally sufficient to protect against an oxidative stress when sequestrases are active. We found that Hsp104 suppressed the high levels of protein aggregation formed in the *btn2 hsp42* mutant following ROS exposure consistent with the idea that the Hsp104 disaggregase normally acts to resolve spatially sorted protein aggregates that are formed following oxidant exposure.

CytoQ and INQ have primarily been characterized using model fluorescently labelled misfolded reporters and little is known regarding the *in vivo* substrates of Btn2 and Hsp42. We focussed on the Sup35 protein since oxidative damage to Sup35 has been shown promote Sup35 aggregation as well as being an important trigger influencing the formation of heritable [*PSI*^+^] prions (Grant, 2015; Jamar *et al*., 2017; Speldewinde *et al*., 2017). Hydrogen peroxide exposure increased the number of Sup35 puncta in cells, and in the absence of Btn2 and Hs42, the number of cells containing Sup35 puncta was further increased with many cells containing multiple Sup35 puncta. The increased Sup35 aggregation detected in *btn2 hsp42* mutants required the Sup35 PrD which has previously been implicated in the formation of non-amyloid Sup35 phase separated aggregates during nutrient depletion stress conditions (Franzmann *et al*., 2018). Our model is that soluble proteins, such as Sup35, undergo misfolding in response to oxidative damage, and the oxidatively damaged proteins are triaged by Btn2 and Hsp42 to the various protein deposit sites in cells (Fig. 7). In the absence of Hsp42 and Btn2, protein sequestration is deficient and non-specific protein aggregates accumulate in cells. These aggregates are targeted by Hsp104 and other chaperones, but non-specific aggregates ultimately overwhelm the protein homeostasis machinery resulting in sensitivity to oxidative stress conditions. This implicates protein sequestration as key antioxidant defence mechanism that functions to mitigate the damaging consequences of protein oxidation and resulting protein misfolding and aggregation.

**Fig. 7.**
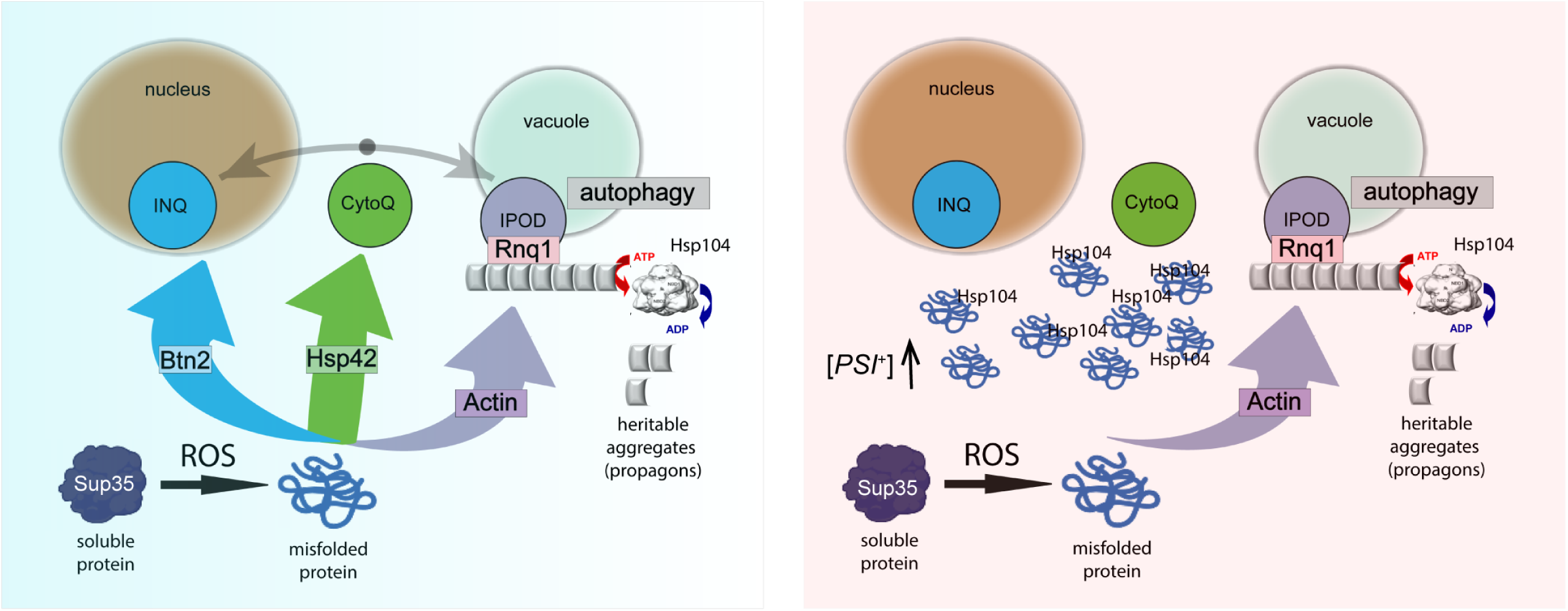
Protein sequestration is a key antioxidant defence mechanism that functions to mitigate the damaging consequences of protein oxidation. Soluble proteins that undergo misfolding in response to oxidative damage are normally triaged by Btn2 and Hsp42 to the various protein deposit sites in cells. In the absence of Hsp42 and Btn2, protein sequestration is deficient and non-specific protein aggregates accumulate in cells. These aggregates are targeted by Hsp104 and other chaperones, but non-specific aggregates ultimately overwhelm the protein homeostasis machinery resulting in sensitivity to oxidative stress conditions. *De novo* prion formation following protein oxidation depends on IPOD localization which acts as a sorting centre determining whether oxidized proteins are cleared via autophagy, or alternatively, form heritable protein aggregates (prions), dependent on Hsp104. Prion formation is increased in *btn2 hsp42* mutants in response to ROS exposure independent of IPOD localization.

[*PSI*^+^] is the amyloid prion form of the Sup35 protein (Wickner, 1994). We found that [*PSI*^+^] prion formation is unaffected in *btn2 hsp42* mutants during normal non-stress growth conditions. There are a plethora of anti-prion systems present in yeast that may be sufficient to maintain the low frequency of [*PSI*^+^] prion formation that occurs during non-stress conditions even in sequestrase mutants (Wickner, 2019). This contrasts with the [*URE3*] prion, where Btn2 and Cur1 normally act to suppress [*URE3*] formation in in a mechanism that requires Hsp42 (Kryndushkin *et al*., 2008; Wickner *et al*, 2014). Btn2 co-localizes with Ure2 and is thought to cure [*URE3*] by sequestering amyloid filaments preventing their inheritance during cell division. Overexpression of Btn2 has also been shown to cure cells of the [*URE3*] prion (Kryndushkin *et al*., 2008). In contrast, overexpression of Btn2 does not cure the [*PSI*^+^] prion, despite Btn2 co-localizing with Sup35 protein aggregates (Barbitoff *et al*, 2017; Kryndushkin *et al*., 2008; Zhao *et al*, 2018). Overexpression of Hsp42 also cures [*URE3*] (Wickner *et al*., 2014) but not [*PSI+*] (Zhao *et al*., 2018). However, another study has shown that the frequency of overexpression-induced [*PSI*^+^] formation is increased in an *hsp42* mutant, and overexpression of Hsp42 effectively cures [*PSI*^+^] suggesting that Hsp42 can protect against [*PSI*^+^] prion formation (Duennwald *et al*, 2012). We found that the number of cells containing Sup35 puncta increased in response to oxidative stress and the colocalization of these puncta with Btn2 and Hsp42 was maintained following hydrogen peroxide exposure. Btn2 and Hsp42 appear to act as an anti-prion system that specifically suppress [*PSI*^+^] formation during ROS exposure, but not during overexpression-induced prion formation. This is consistent with the idea that Btn2 and Hsp42 function to sequester oxidatively damaged proteins, thus preventing their templating to form the heritable prion form.

The IPOD is a site of accumulation of amyloidogenic proteins, as well as oxidatively damaged proteins (Kaganovich *et al*., 2008; Rothe *et al*., 2018; Tyedmers *et al*., 2010). *De novo* prion formation following protein oxidation depends on IPOD localization which acts as a sorting centre determining whether oxidized proteins are cleared via autophagy, or alternatively, form heritable protein aggregates (prions), dependent on Hsp104 (Fig. 7). IPOD-like inclusions have also been identified in mammalian cells, although little is known regarding their functional significance (Hipp *et al*, 2012; Kaganovich *et al*., 2008; Weisberg *et al*, 2012). One possibility is that proteins that are not degraded or re-solubilized at CytoQ/INQ may be trafficked to the IPOD which is thought to harbour terminally misfolded proteins (Escusa-Toret *et al*., 2013; Kaganovich *et al*., 2008). Hsp42 and Btn2 are thought to play roles in targeting terminally misfolded proteins including amyloidogenic substrates to the IPOD (Kaganovich *et al*., 2008; Rothe *et al*., 2018; Tyedmers *et al*., 2010). The resulting localised concentration of these proteins can facilitate the nucleation of prion protein polymerisation. This involves a two-stage process which initially involves the formation of non-transmissible extended polymers of the prion protein, followed by their fragmentation into shorter transmissible propagons, catalysed by Hsp104 and similar chaperones (Tyedmers *et al*., 2010). The elevated frequency of oxidant-induced [*PSI*^+^] prion formation in *btn2 hsp42* mutants did not require IPOD localisation since it was unaffected in cortical actin mutants which have previously been shown to disrupt Sup35 IPOD localisation (Speldewinde *et al*., 2017). This raises the question as to how the frequency of oxidant-induced [*PSI*^+^] prion formation is elevated in sequestrase mutants. We suggest there are two possibilities by which this may occur. Firstly, the high localized concentrations of Sup35 at multiple non-specific aggregation sites may increase the likelihood of prion induction. For example, other aggregating glutamine (Q)/asparagine (N)-rich proteins can promote *de novo* [*PSI*^+^] prion formation (Arslan *et al*., 2015). Secondly, chaperones and other anti-prion systems may become overwhelmed by the high levels of non-specific protein aggregates present in *btn2 hsp42* mutants meaning they are not available to suppress [*PSI*^+^] prion formation. These two possibilities are not mutually exclusive and are analogous to the cross-seeding and titration models that have been proposed to explain the requirement for [*PIN*^+^] in *de novo* induction of [*PSI*^+^] formation (Derkatch & Liebman, 2007; Keefer *et al*, 2017).

## MATERIALS AND METHODS

### Yeast strains and plasmids

All yeast strains used in this study were derived from 74D-694 *(MATa ade1-14 ura3-52 leu2-3,112 trp1-289 his3-200)*. Strains were deleted for *BTN2* (*btn2::TRP1, btn2::LEU2*)*, HSP42* (*hsp42::TRP1*)*, CUR1* (*cur1::LEU2*) and *ABP1* (*abp1::loxLE-hphNT1-loxRE*) using standard yeast methodology. Btn2 and Hsp42 were C-terminally Myc or GFP tagged using a PCR-based approach (Knop *et al*, 1999). Yeast strains expressing *SUP35-GFP* or *ΔN-SUP35-GFP* were constructed using a plasmid shuffle approach (Parham *et al*, 2001). Briefly, a yeast strain deleted for the chromosomal copy of SUP35 was complemented with a *URA3-CEN* plasmid carrying the wild-type *SUP35* gene. *SUP35-GFP* and *ΔN-SUP35-GFP* were constructed by cloning commercially synthesized gene fragments into plasmid pRS413 (Sikorski & Hieter, 1989). 5-Fluoro-orotic acid (5-FOA)-containing medium was used to select for cells expressing GFP-tagged versions of Sup35. The yeast plasmid *CUP1-SUP35NM-GFP* expressing the Sup35NM domain conjugated to RFP under the control of the *CUP1* promoter has been described previously (Patino *et al*., 1995) as have the yeast plasmids expressing *CUP1-RNQ1-CFP, HSP104-*RFP and *GDP-HSP104*-mCherry (Lee *et al*., 2010; Malinovska *et al*., 2012; Sondheimer & Lindquist, 2000).

### Growth and stress conditions

Yeast strains were grown at 30°C with shaking at 180 rpm in minimal SD media (0.67% w/v yeast nitrogen base without amino acids, 2% w/v glucose) supplemented with appropriate amino acids and bases. Stress sensitivity was determined by growing cells to exponential phase in SD media and spotting diluted cultures (A_600_ = 1.0, 0.1, 0.01, 0.001) onto SD agar plates containing various concentrations of hydrogen peroxide. Respiratory growth media contained 3% (w/v) glycerol and 1% v/v ethanol instead of glucose. For oxidative stress conditions, cells were grown to exponential phase in SD media and treated with 0.8 mM hydrogen peroxide for one hour.

### Protein and Western blot analysis

Protein extracts were electrophoresed under reducing conditions on SDS-PAGE minigels and electroblotted onto PVDF membrane (Amersham Pharmacia Biotech). Primary antibodies used were raised against Sup35 (Ness *et al*, 2002), Rnq1 (Sideri *et al*., 2011), Myc (Myc 4A6, Millipore), GFP (Invitrogen), Pgk1 (ThermoFisher Scientific), DNPH (Dako) and Hsp104 (Abcam). The analysis of Sup35 amyloid polymers by semi-denaturing detergent-agarose gel electrophoresis (SDD-AGE) was performed as described previously (Alberti *et al*, 2010). Protein carbonylation was measured by reacting carbonyl groups with 2,4-dinitrophenyl-hydrazine (DNPH) and detected using rabbit anti-DNPH (Shacter *et al*, 1994). Insoluble protein aggregates were isolated as previously described and visualized by silver staining (Hamdan *et al*., 2017).

### Analysis of prion formation

Prion formation was quantified based on readthrough of the nonsense (UGA) mutation in the *ADE1* gene as described previously (Speldewinde *et al*., 2017). [*psi*^−^] strains are auxotrophic for adenine and appear red due to the accumulation of an intermediate in the adenine biosynthesis pathway, whereas, [*PSI*^+^] strains give rise to white/pink Ade^+^ colonies due to suppression of the *ade1-14* nonsense mutation and production of functional Ade1 protein. Diluted cell cultures were plated onto SD plates lacking adenine (SD-Ade) and incubated for 7-10 days. Prions were differentiated from nuclear gene mutations by their irreversible elimination on plates containing 4mM GdnHCl. GdnHCl effectively blocks the propagation of yeast prions by inhibiting the key ATPase activity of Hsp104, a molecular chaperone that functions as a disaggregase and is required for prion propagation (Ferreira *et al*, 2001; Jung & Masison, 2001). Data shown the means of at least three independent biological repeat experiments expressed as the number of [*PSI*^+^] cells relative to viable cells. Data are presented as mean values ± SD. Statistical analysis for multiple groups was performed using one-way ANOVA.

### Fluorescence microscopy

Cells were harvested, resuspended in deionized water, and spotted onto poly-L-lysine microscopy slides. Cells were imaged with a z-spacing of 0.2 µm using a Leica 100x/1.40-0.70 NA Oil Plan objective lens and K5 cMOS camera fitted onto the Leica DM550 B microscope (LEICA Microsystems GmbH; Wetzlar, Germany). Image acquisition was supported by the Leica Application Suite X (LAS X v.3. 7.4.23463) and processing was conducted using lmageJ.

## Acknowledgements.

We thank Lynn Megeney, Mick Tuite and Simon Alberti for generously providing plasmids and antibodies used in the study.

## Author contributions

ZC, KK, DRC and CMG performed the experiments and analysed the data. CMG wrote the manuscript, designed and supervised the project.

